# Development of the branchial musculature of the Siberian sturgeon (*Acipenser baerii*) reveals a heterochronic shift during the evolution of acipenseriform cranial muscles

**DOI:** 10.1101/2023.02.14.528484

**Authors:** Benjamin Naumann, Peter Warth, Jörg U. Hammel, Julian Moosmann, Peter Konstantinidis, Lennart Olsson

## Abstract

Heterochronic shifts are regarded one of the major evolutionary changes acting on developmental modules and underlying the origin of morphological disparity. Conserved characters, rarely subject to heterochronic shifts during the curse of evolution, in contrast could indicate underlying developmental or functional constraints. Here we use the development of the cranial musculature Siberian sturgeon (*Acipenser baerii*) as a model to investigate the role of heterochrony during the evolution of the craniofacial system of Actinopterygii. Using histology, fluorescent antibody staining and fast propagation-based phase contrast imaging in combination with 3D-reconstruction we describe the development of the branchial and hypobranchial musculature. We show that the development of the first branchial arch is accelerated compared to other basal-branching actinopterygians leading to a more synchronous development with the hyoid arch. A pattern that could relate to the derived migratory behaviour of the neural crest cells in sturgeons. In contrast, the developmental timing of the more posterior branchial musculature, including the *cucullaris* muscle in the Siberian sturgeon, appears to be highly conserved compared to other Actinopterygii and even Osteognathostomata. This could indicate the presence of functional or developmental constraints underlying the evolution of the muscles at the head/trunk interface.

## 1 Introduction

Heterochrony refers to phylogenetic differences in developmental rate and timing and is regarded as a major evolutionary mechanism underlying morphological disparity (Gould, 1977; Hall, 2012; Mabee, Olmstead, & Cubbage, 2000). While some morphological characters exhibit large-scale heterochronic shifts (developmental acceleration or delay), others may vary only slightly or not at all when compared between different taxa. Such conserved developmental rates and timing might result from functional and developmental constraints limiting the evolvability (*sensu* Arthur, 2021) of these characters (Gould & Lewontin, 1979). The craniofacial musculature and skeleton of vertebrates has been proven as a valuable system to investigate the role of heterochronic shifts in the evolution of morphological disparity (Camacho et al., 2020; Mitgutsch, Piekarski, Olsson, & Haas, 2008; Morris, Vliet, Abzhanov, & Pierce, 2019; Naumann, Warth, Olsson, & Konstantinidis, 2017b; Tokita, Kiyoshi, & Armstrong, 2007). With over 36000 species, ray-finned fishes (Actinopterygii) account for over half of the extant vertebrate species (Fricke, Eschmeyer, & Van der Laan, 2018) (Figure 1a). It is assumed that the main driver of this evolutionary success (in terms of species numbers) is their diverse craniofacial morphology resulting in a high diversity of feeding mechanisms (Lauder, 1982; Warth, Hilton, Naumann, Olsson, & Konstantinidis, 2018). While the cranial morphology of many actinopterygian species is well described, the developmental patterns underlying the observed disparity is less well studied. Therefore, studies on the craniofacial development of various actinopterygians still represent an excellent framework to ask the question: What are the roles of developmental constraints and heterochronic shifts in the evolution of morphological disparity?

**Figure 1.**
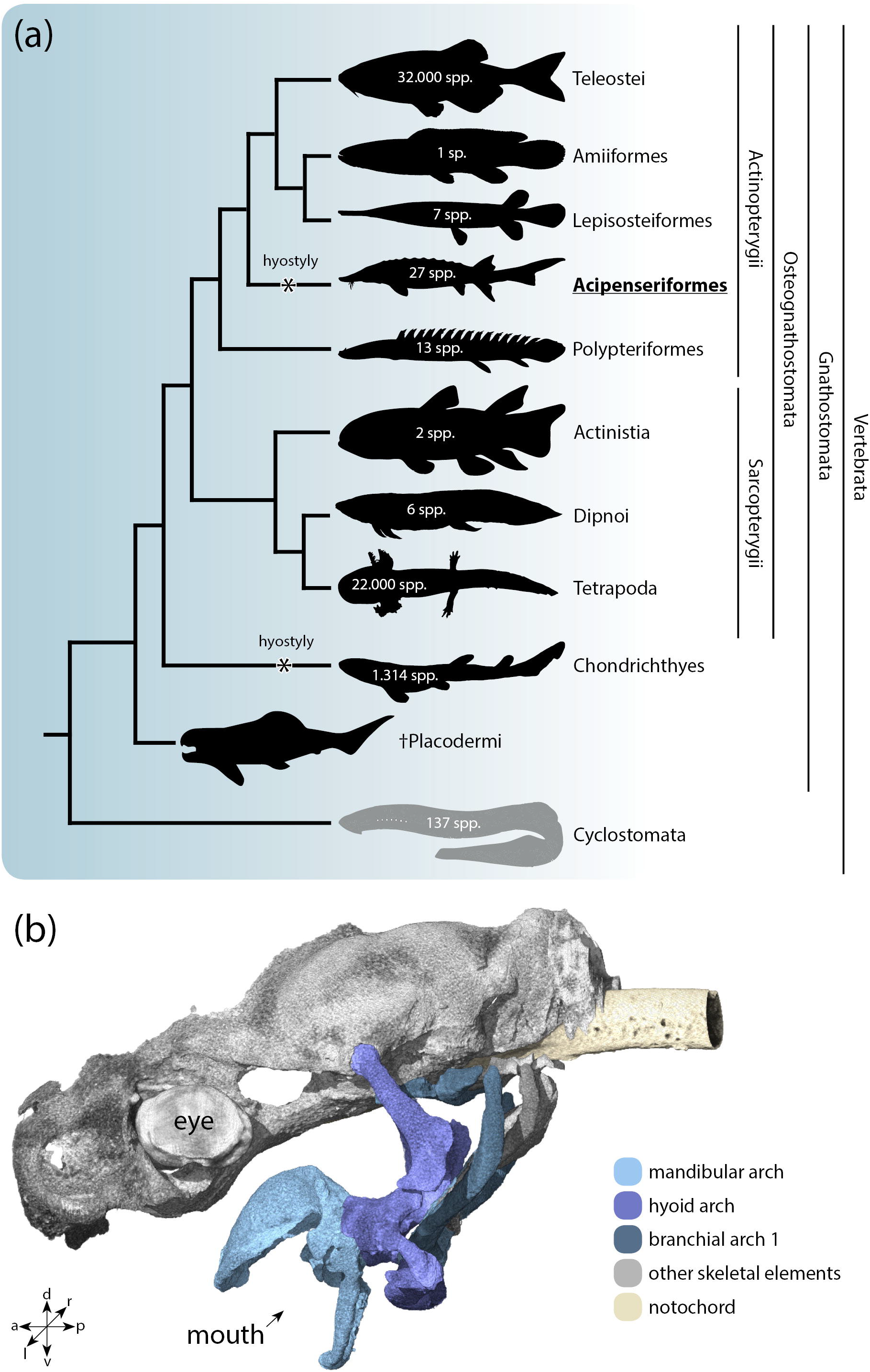
(a) Cladogram of extant vertebrates. Numbers within the pictograms represent the approximate number of extant species (Froese & Pauly, 2022). A hyostylic jaw suspension independently evolved (black asterisk) in the stem lineage of Acipenseriformes (sturgeons) and within Chondrichthyes. The grey color in Cyclostomata indicates the absence of a *cucullaris*. The presence of a *cucullaris* has been proposed in species of the extinct Placodermi (Trinajstic et al., 2013). (b) 3D reconstruction of the neuro- and viscerocranium of a juvenile *Acipenser baerii* at stage 45.

Among early-branching ray-finned fishes, sturgeons and paddle-fishes (Acipenseriformes) exhibit a highly derived craniofacial morphology. It is characterized by a partial retention of a relatively undifferentiated, “embryonic” configuration of the musculoskeletal system with delayed development or complete absence of endoskeletal ossifications present in other actinopterygian lineages (Carroll & Wainwright, 2003). Additionally, sturgeons share a uniquely derived mode of jaw suspension (Bemis et al., 1997; Miller, 2005). The mandibular skeleton is not connected to the neurocranium but suspended via hyoid skeletal elements (Gregory, 1904) (Figure 1b). This specific hyostylic condition is highly derived, yet in some sharks and batoid skates (Chondrichthyes), a convergent jaw suspension has evolved independently (Wilga, 2005). In both, sturgeons and some chondrichthyan groups, this condition allows an extreme protrusion of the upper jaw and plays a crucial role in suction feeding (Carroll & Wainwright, 2003). Associated with this hyostylic condition in sturgeons, the musculature of the mandibular and hyoid arch is also derived compared to other actinopterygians (Arratia & Schultze, 1991; Warth, Hilton, Naumann, Olsson, & Konstantinidis, 2017), yet secondarily simplified (Edgeworth, 1935) by developmental truncation. Paedomorphosis, a heterochronic shift resulting in delayed onset of development and early truncation of musculoskeletal differentiation, is thought to play a major role in establishing this morphological condition in sturgeons (Bemis, Findeis, & Grande, 1997; Tsessarsky, 2020, 2022; Warth et al., 2017, 2018).

Here we investigate the development of the branchial and hypobranchial musculature in the Siberian sturgeon (*Acipenser baerii*). This study complements our previous studies on craniofacial development of non-teleostean actinopterygians (Konstantinidis et al., 2015; Naumann, Warth, Olsson, & Konstantinidis, 2017a; Warth et al., 2017, 2018). We show that the muscle development of the first branchial arch is accelerated compared to the more posterior branchial arches leading to a more synchronous development of the muscles of the hyoid and first branchial arch. This indicates a heterochronic shift compared to other actinopterygians. The remaining branchial arch muscles develop in a conserved anterior-to-posterior sequence. In contrast, the developmental timing of the posterior-most branchial arch muscles and the cucullaris muscle seem to be conserved compared to other ray-finned (Diogo, Hinits, & Hughes, 2008; Edgeworth, 1935; Naumann et al., 2017a; Noda, Miyake, & Okabe, 2017) and lobe-finned taxa (Edgeworth, 1935; Naumann & Olsson, 2018; Theis, 2010; Theis et al., 2010; Ziermann, Clement, Ericsson, & Olsson, 2018; Ziermann & Diogo, 2013). This might indicate developmental constraints (e.g., based on migratory patterns of neural crest cells) limiting deviations from an ancestral pattern for the *cucullaris* and other muscles developing at the head-trunk interface.

## 2 Materials and Methods

### 2.1 Specimens

Specimens of the Siberian sturgeon (*Acipenser baerii* Brandt 1869) were obtained from artificial spawning at the “Fischzucht Röhnforelle” (Gersfeld, Germany). Embryos and larvae were raised and treated as described previously (Warth et al., 2018). Specimens were staged according to the staging tables of *Acipenser gueldenstaedtii* Brandt & Ratzeburg 1833 (Ginsburg & Dettlaff, 1991; Schmalhausen, 1991). A specimen list is given in Supplementary Material 1.

### 2.2 Terminology

For the terminology of the musculature we follow mainly the terminology used for Teleostei (Winterbottom, 1973) with one exception: forthe muscle originating on the lateral surface of the otic capsule and inserting on the supracleithrum, we use the term *cucullaris* according to Edgeworth (1935) instead of protractor pectoralis (Winterbottom, 1973) to facilitate comparison with tetrapods.

### 2.3 Histology

Specimens fixed in 4 % saline-buffered paraformaldehyde (PFA) were embedded in paraffin and sectioned into 7 μm transverse sections using a HM360 microtome (Microm). Sections were stained using the Resourcin-Trichrome and Heidenhain’s AZAN technique (Böck, 1989). Single sections were digitized with an Olympus BX51 microscope using the dotslide software 2.3 (Olympus Corporation).

### 2.4 Fluorescent whole mount antibody staining

Specimens were fixed and stored in Dent’s fixative (four parts 100% methanol, one part dimethyl sulfoxide) or PFA. Subsequent whole mount antibody staining was performed as described by Warth et al. 2018. Anti-desmin (1/500, Monosan, PS031), anti-newt skeletal muscle (1/100, DSHB, 12/101), anti-acetylated alpha-tubulin (1/500, Sigma, T6793) and anti-collagen type II (1/100, DSHB, CIIC1) were used as primary antibodies. Alexa488-anti-mouse (1/500, Thermo Fisher Scientific, #R37120) and Alexa568-anti-rabbit (1/500, Thermo Fisher Scientific, #A-11011) were used as secondary antibodies. Digital stacks from the antibody staining were obtained using a Zeiss LSM 510 confocal microscope and processed with the ZEN 2009 and 2012 Software (Carl Zeiss).

### 2.5 Propagation-based phase contrast imaging

Specimens were fixed in 4 % saline-buffered paraformaldehyde (PFA) and subsequently rinsed and transferred in a series of ascending alcohol concentrations (30%, 50%, 70%, 90%, 99%) before immersion in an ethanol based 1 % iodine solution. No particular emphasis was put on the time of immersion or the subsequent wash steps before imaging. However, the minimum time of incubation was 24 hours for the smallest and 1 week for the largest specimens. The washing step took 15 minutes minimum for the smallest and 30 minutes minimum for the largest specimens. Experiments were performed at the P05 Imaging Beamline (Greving et al., 2014; Haibel et al., 2010; Wilde et al., 2016), which is operated by Helmholtz–Zentrum Hereon at PETRA III at Deutsches Elektronen-Synchrotron (DESY) in Hamburg, Germany, both members of the Helmholtz Association HGF. X-rays were monochromatized to a photon energy of 35 keV using a double crystal monochromator. An indirect detector system was used comprised of a 20 MPixel CMOS camera with an effective pixel size of 0.64 μm, developed by HEREON and the Karlsruhe Institute of Technology (Lautner et al., 2017; Lytaev et al., 2014), a 100 μm thick CdWO4 scintillator and a 10 × microscope optics. The tomograms were reconstructed using a routine (Moosmann, 2021; Moosmann et al., 2014) implemented in MATLAB using the ASTRA toolbox for tomographic backprojection (Palenstijn, Batenburg, & Sijbers, 2011; van Aarle et al., 2016; van Aarle et al., 2015) resulting in an effective voxel size of 1.28 μm in the reconstructed tomographic volume after two times binning of raw data.

### 2.6 Image processing, segmentation and 3D rendering

Image adjustments were conducted using Fiji (Schindelin et al., 2012) or Adobe Photoshop CS6. 3D reconstructions based on the digital image stacks were performed semi-manually using Amira 5.2 3D-analysis software (FEI Visualization Sciences Group) and transferred to VG Studio Max 2.0 (Volume Graphics) for volume renderings.

## 3 Results

### 3.1 The anatomy and innervation of the branchial and hypobranchial muscles

The innervation (I), origin (o), course (c) and insertion (i) are listed for every muscle based on antibody staining and a μCT-scanning of juveniles (based on stages 44 to 46) and dissection of a larger juvenile (total length: 19.5 cm). The anatomy of the muscles is shown in Figure 2, the innervation pattern in Figure 3. Skeletal elements are shown in Supplementary Material 2 according to Warth et al. (2017).

**Figure 2.**
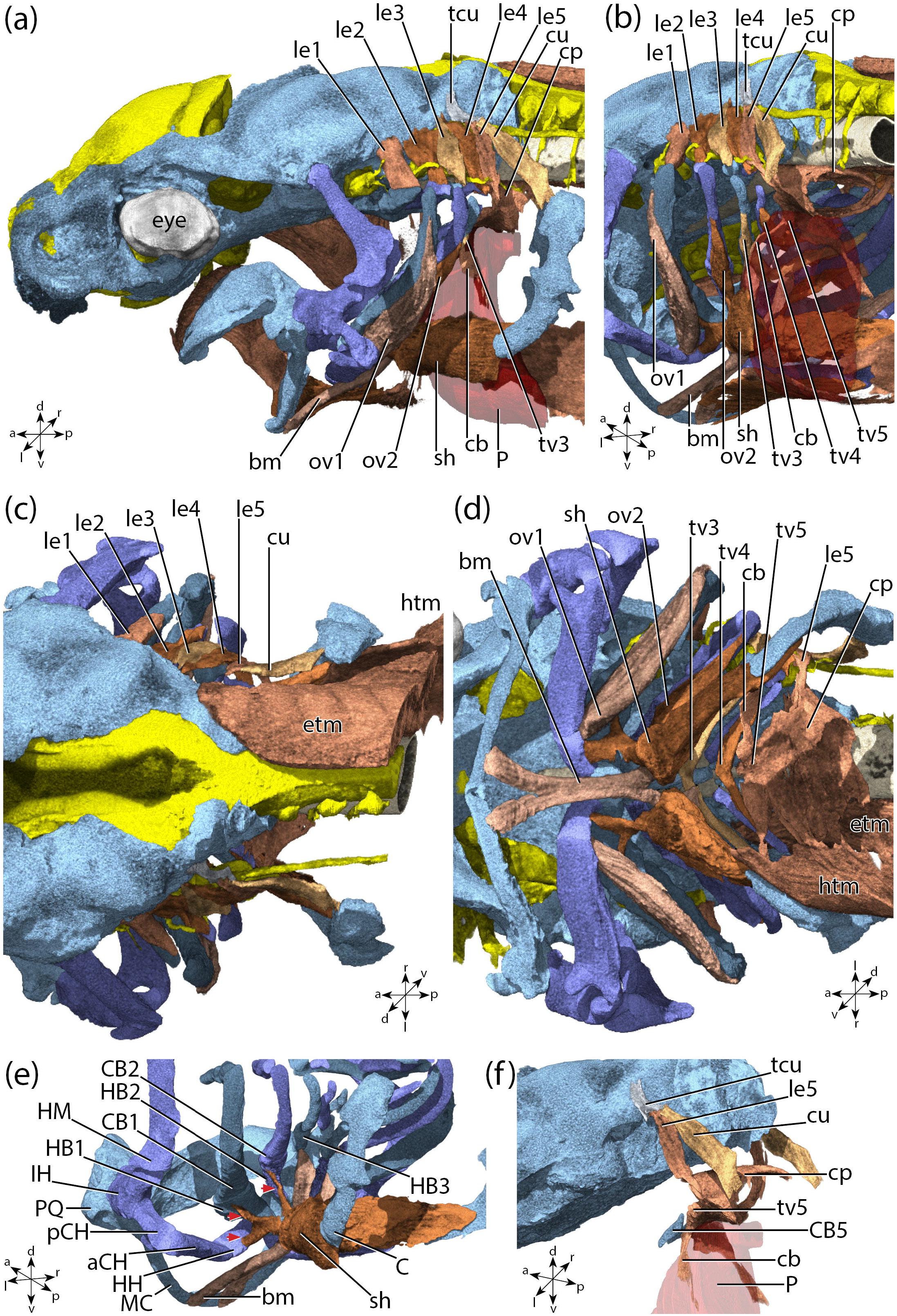
3D rendering of the segmented tomogram of the skeleton (shades of blue), muscles (shades of red) and nerves (shades of yellow) of the head of a juvenile *Acipenser baerii* at stage 45. (a) lateral view. (b) posterolateral view. The shoulder girdle has been removed. (c) dorsal view. (d) ventral view. The pericard has been removed. (e) posterolateral view of the ventral branchial and hypobranchial muscles. Red arrows indicate the insertion sited of the sternohyoideus muscle. (f) lateral view of the muscles of branchial arch 5. Skeletal elements of branchial arch 1 to 4 have been removed.

**Figure 3.**
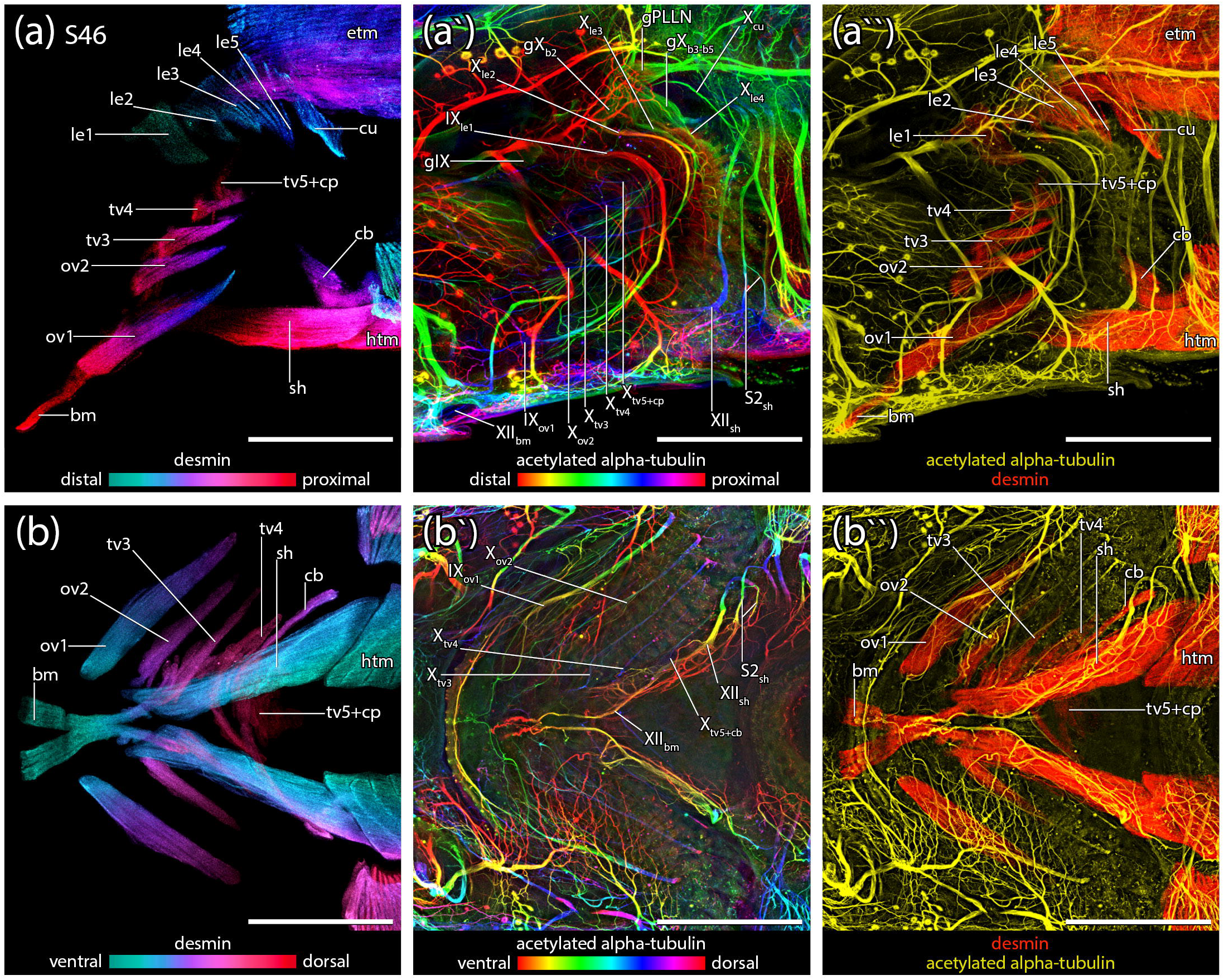
Maximum intensity projections of a whole-mount antibody staining against desmin and acetylated alpha-tubulin showing branchial muscles and their innervation of a *Acipenser baerii* juvenile at stage 46. Upper row (a to a``), posterior branchial region from lateral view; (a) maximum intensity projection of an antibody staining against desmin. Colors are depth-coded indicated by the color scale. (a`) maximum intensity projection of an antibody staining against acetylated alpha-tubulin. Colors are depth-coded indicated by the color scale. (a) merged maximum intensity projection of an antibody staining against desmin and acetylated alpha-tubulin. Muscles are colored in red (desmin), nerves in yellow (acetylated alpha-tubulin). (b to b``) posterior branchial region from ventral view. (a) maximum intensity projection of an antibody staining against desmin. Colors are depth-coded indicated by the color scale. (a`) maximum intensity projection of an antibody staining against acetylated alpha-tubulin. (a) merged maximum intensity projection of an antibody staining against desmin and acetylated alpha-tubulin. Muscles are colored in red (desmin), nerves in yellow (acetylated alpha-tubulin). Scale bar is 500 μm.

#### Branchial arch 1

***Levator externus* 1**: (I) dorsal ramulus of posttrematic branchial ramus of the glossopharyngeal nerve, (o) lateral surface of the otic capsule, (c) lateroventrad, (i) dorsal surface of epibranchial 1; ***Obliquus ventralis* 1**: (I) ventral ramuli of posttrematic branchial ramus of the glossopharyngeal nerve; (o) hypohyal and anterior ceratohyal, spanning the joint between these two cartilages, (c) posterodorsad, (i) ceratobranchial 1.

#### Branchial arch 2

***Levator externus* 2**: (I) dorsal ramulus of the posttrematic branchial ramus of the vagal nerve arising from the anterior vagal ganglion, (o) lateral surface of the otic capsule, posterior to the *levator externus* 1, (c) lateroventrad, (i) dorsal surface of epibranchial 2; ***Obliquus ventralis* 2**: (I) ventral ramulus of the posttrematic branchial ramus of the vagal nerve arising from the anterior vagal ganglion, (o) hypobranchial 2, (c) posterodorsad, (i) ceratobranchial 2.

#### Branchial arch 3

***Levator externus* 3**: (I) dorsal ramulus of the posttrematic branchial ramus of the vagal nerve arising from the posterior vagal ganglion, (o) lateral surface of the otic capsule, posterior to the *levator externus* 2, (c) lateroventrad, (i) dorsal surface of epibranchial 3; ***Transversus ventralis* 3**: (I) ventral ramulus of the posttrematic branchial ramus of the vagal nerve arising from the posterior vagal ganglion, (o) median raphe with its antimere; (c) posterodorsad, (i) ceratobranchial 3.

#### Branchial arch 4

***Levator externus* 4**: (I) dorsal ramulus of the posttrematic branchial ramus of the vagal nerve arising from the posterior vagal ganglion, (o) lateral surface of the otic capsule, posterior to the *levator externus* 3 and closely connected to the *levator externus* 5, (c) lateroventrad, (i) dorsal surface of epibranchial 4; ***Transversus ventralis* 4**: (I) ventral ramulus of the posttrematic branchial ramus of the vagal nerve arising from the posterior vagal ganglion, (o) median raphe with its antimere, (c) posterodorsad, (i) ceratobranchial 4.

#### Branchial arch 5 and posterior

***Levator externus* 5**: (I) dorsal ramulus of the posttrematic branchial ramus of the vagal nerve arising from the posterior vagal ganglion, (o) lateral surface of the otic capsule from a common fascia with the *cucullaris*, posterior to the *levator externus* 4 (c) lateroventrad around the pharynx, (i) tapers into the common muscle mass of the *constrictor pharyngeus*, *transversus ventralis* 5 and *coracobranchialis*; ***Cucullaris***: (I) distinct dorsal vagal ramus, originating from the posterior vagal ganglion; (o) lateral surface of the otic capsule from a common fascia with the *levator externus* 5, (c) posteroventrad, (i) anterodorsal surface of the suprascapula; ***Transversus ventralis* 5**: (I) ventral ramulus of the posttrematic branchial ramus of the vagal nerve arising from the posterior vagal ganglion, also innervating the *constrictor pharyngeus* and *coracobranchialis*, (o) median raphe with its antimere, (c) posterodorsad, (i) tapers into the *constrictor pharyngeus*; ***Constrictor pharyngeus***: (I) ventral ramulus of the posttrematic branchial ramus of the vagal nerve arising from the posterior vagal ganglion, also innervating the *transversus ventralis* 5 and *coracobranchialis*, (o,i) pharyngeal wall, (c) encircles the pharyngeal wall at the level of the posterior half of the ceratobranchial 5, ventrally in close contact with the pericardial wall. **C***oracobranchialis*: (I) ventral ramulus of the posttrematic branchial ramus of the vagal nerve arising from the posterior vagal ganglion, also innervating the *transversus ventralis* 5 and *constrictor pharyngeus*, (o) a common muscle mass together with the *constrictor pharyngeus* and the *transversus ventralis* 5, (c) ventrad closely connected to the pericardial wall, (i) anterodorsal margin of the coracoid.

#### Hypobranchial muscles

***Sternohyoideus***: (I) spinal hypoglassal nerve, posterior end also innervated by some ramuli of the second spinal nerve, (o) anterior edge of the coracoid, (c) anterad, (i) three thin tendons that insert on the hypobranchial 2, 1 and the hypohyal each; ***Branchiomandibularis***: (I) anterior ramuli of the hypoglossal nerve branching off from a common ramus together with the *sternohyoideus*, (o) hypobranchial 3, (c) anterad, (i) posterior surface of Meckel`s cartilage.

### 3.2 The development of the branchial and hypobranchial muscles

The complete development of the branchial and hyobranchial muscles is summarized in Table 1.

**Table 1.**
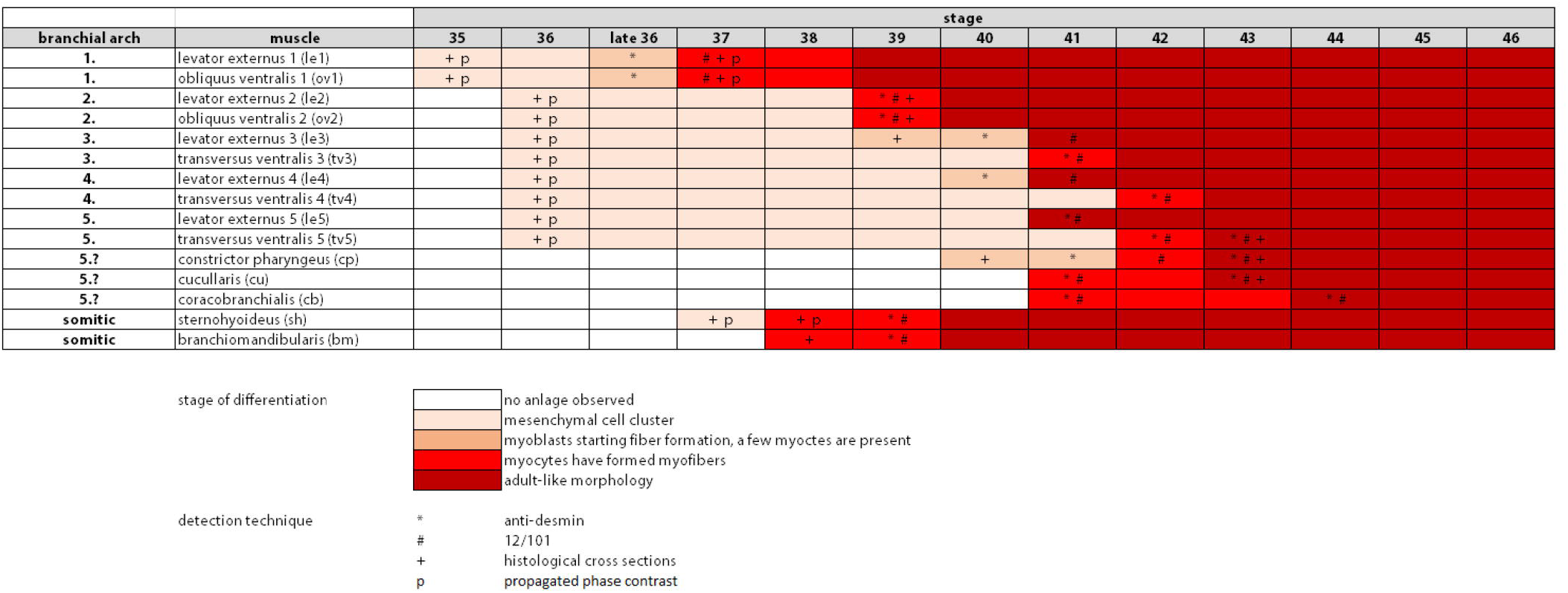
Summary of the development of the branchial and hypobranchial muscles of *Acipenser baerii*.

**Stage 36**(Figures 4a, b, e and 5a and a`): The anlagen of the branchial arch muscles form a dense, histologically undifferentiated mesenchyme posterior to the yolk-rich endoderm of their corresponding branchial arch. Desmin antibody signals, interpreted as the differentiating anlagen of the *levator externus* 1 and *obliquus ventralis* 1, are detectable at late stage 36.

**Figure 4.**
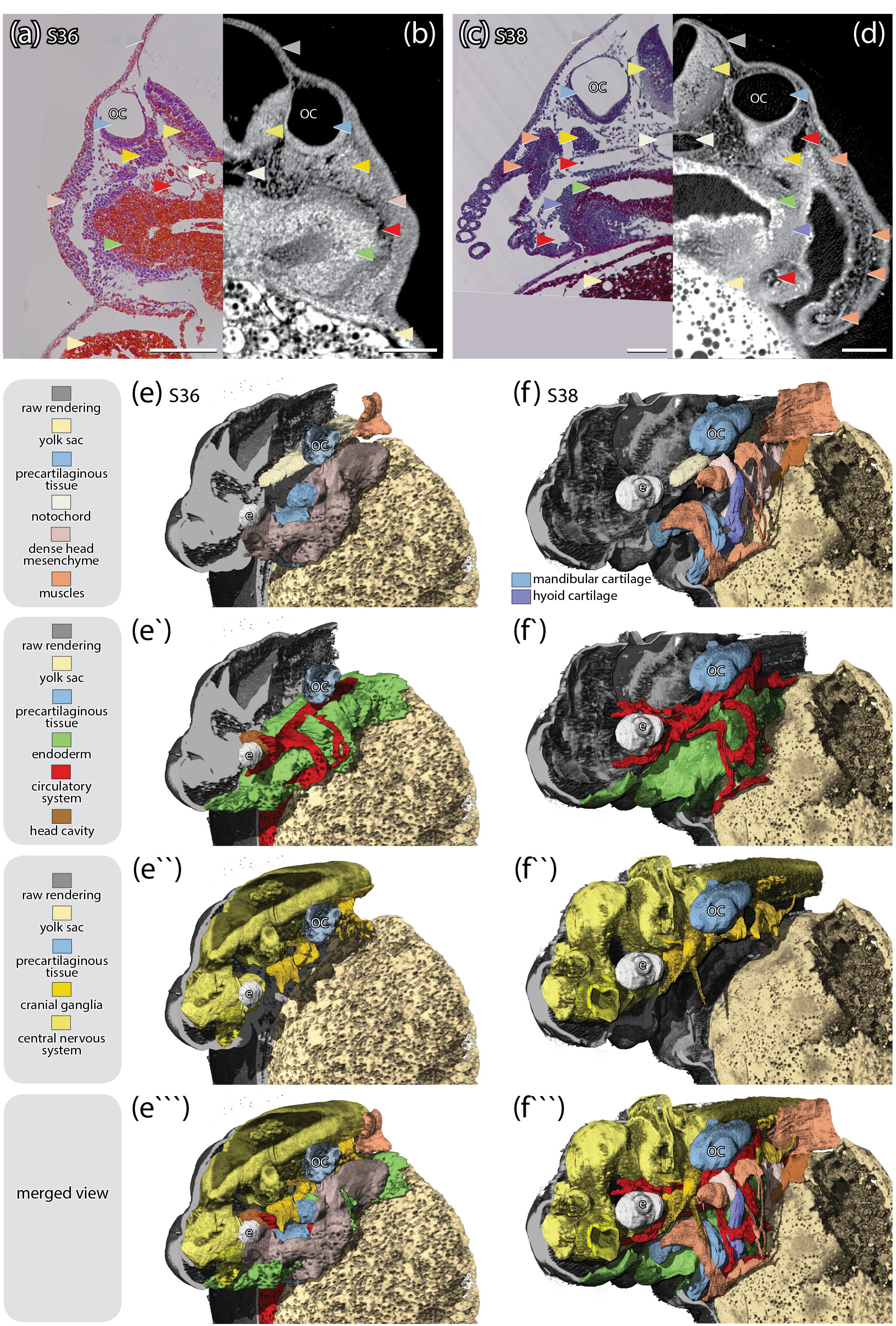
Comparison of Azan-stained histological cross sections and propagation-based phase contrast tomography of *Acipenser baerii* embryos at different developmental stages. Histological cross sections of (a) stage 36 and (c) stage 38 correspond to (b and d) single images of propagated phase contrast scans in region and stage of specimen respectively. (a and c) Colored arrow heads indicate to different anatomical structures and correspond to the colors of the reconstructed organs in (e) to (f```). (e to f```) 3D reconstruction of head region of the specimens shown in (b) and (d) from frontolateral view. (e and f) musculoskeletal system. Cartilage, different shades of blue; eye, white; muscles, different shades of flesh; notochord, ivory; skin, grey; yolk sac, light ochre. (e` and f`) circulatory and digestive system. Gut, green; blood vessels, red. (e`` and f``) nervous system. Brain, light yellow; cranial ganglia, dark yellow. (e``` and f```) merged view. Scale bar is 200 μm.

**Figure 5.**
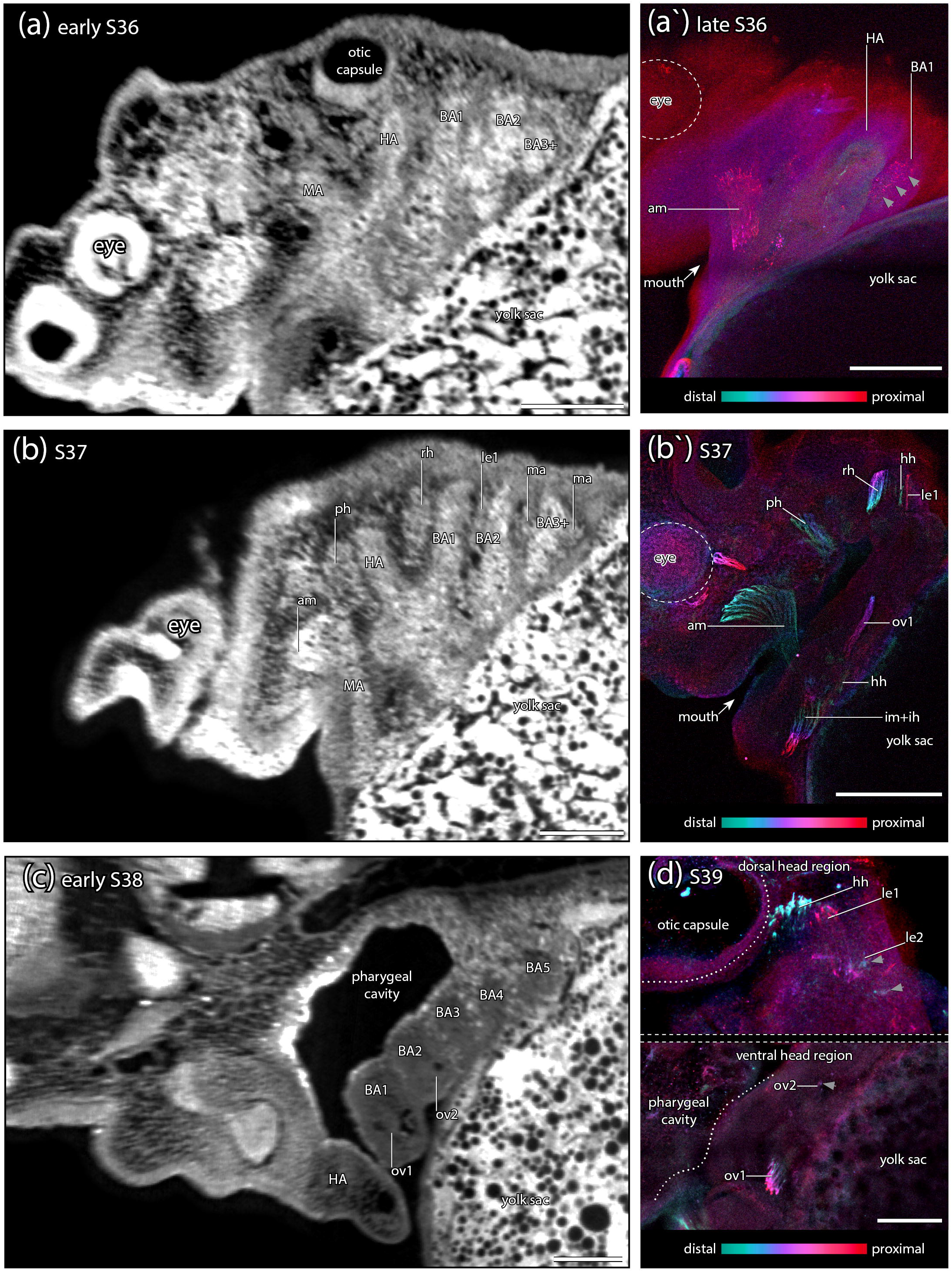
(a to c) single images of propagation-based phase contrast tomography scans of *A. baerii* embryos showing the posterior branchial region in lateral view. (a) early stage 36 (same specimen as reconstructed in Figure 7). (b) stage 37. (c) early stage 38 (same specimen as reconstructed in Figure 7). (a`, b` and d) maximum intensity projection of an antibody staining against desmin. Images show similar regions of the posterior branchial arches as a) to c). grey arrows indicate very weak antibody signals. (d) stage 39. Scale bar is 200 μm.

**Stages 37 and 38**(Figures 4c, d, f and 5b, b` and c): Dense cell aggregations interpreted as muscle primordia are clearly visible in the tomography reconstructions posterior to the pouches of the mandibular, hyoid and branchial arches. An advanced differentiation of the first branchial arch, compared to posterior branchial arches is marked by a clearly separated primordium of the *levator externus* 1 (Figure 5b) The *levator externus* 1 and the *obliquus ventralis* 1 exhibit more distinct desmin and 12/101 antibody signals. While the antibody signal of the *levator externus* 1 is still weak, the *obliquus ventralis* 1 exhibits a very intense signal and has already formed distinct muscle fibers. The comparison between Figure 5b and b` shows, that the actual cell mass of the *levator externus* 1 is much larger as estimated from the desmin antibody signal. The anlagen of the more posterior branchial muscles still appear as a densely packed, histologically undifferentiated mesenchyme.

**Stage 39**(Figure 5d): The *levator externus* 1 and *obliquus ventralis* 1 have increased in size and are both attached to their corresponding skeletal elements. First weak desmin antibody signals, interpreted as the anlagen of the *levator externus* 2 and *obliquus ventralis* 2, are detectable in the second branchial arch.

**Stage 40**(Figure 6a-a``): The *levator externus* 2 and *obliquus ventralis* 2 have increased in size and are both attached to their corresponding skeletal elements. The more posterior dorsal branchial muscles are present as a common mesenchymal mass showing an anterior to posterior differentiation gradient. Anteriorly, the anlage of the *levator externus* 3 forms a long extension exhibiting an intense desmin antibody signal. It extends ventrad into the direction of the developing skeletal elements of the third branchial arch. Posteriorly, the anlage of the *levator externus* 4 forms a shorter cellular band, extending ventrad and exhibiting a weaker desmin antibody signal. The more posterior area of this common muscle anlage is composed of histologically undifferentiated mesenchymal cells exhibiting no desmin antibody signal. Ventrally, the anlagen of the *transversus ventrales* 3 to 5 are present as a compact mesenchyme but have not yet formed muscle fibers. The anlage of the *sternohyoideus* muscle (the hypobranchial chord) is composed of four myotomal portion exhibiting desmin and 12/101 positive muscle fibers.

**Figure 6.**
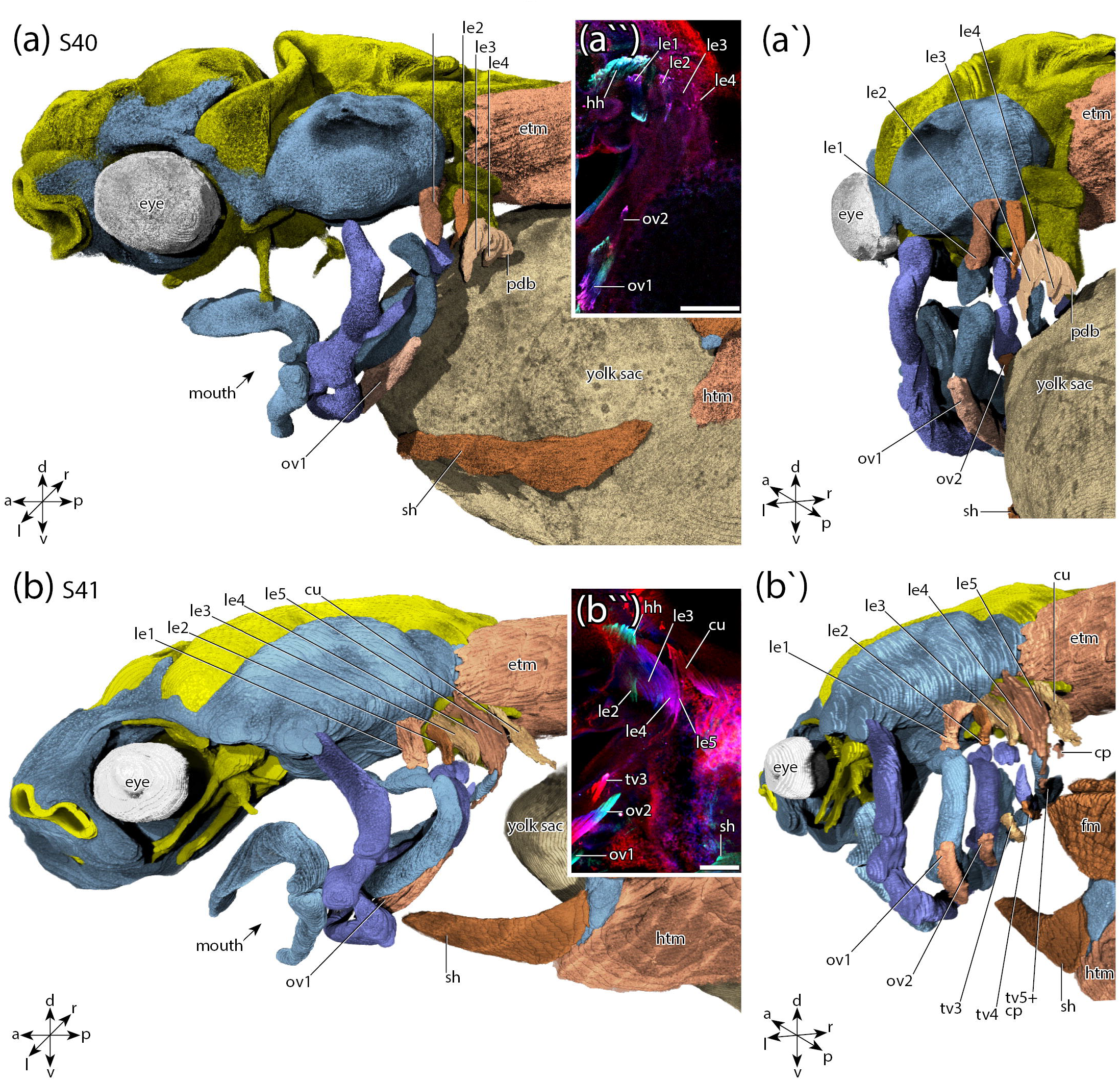
(a to b``) 3D reconstruction of the anatomy of the branchial and hypobranchial musculature in *Acipenser baerii* at different developmental stages. Reconstructions are based on digital image stacks of whole-mount antibody stainings against desmin. (a) stage 40 from lateral view. (a`) posterolateral view. (b) stage 41 from lateral view. dorsal view. (b`) posterolateral view. (a`` and b``) maximum intensity projection of an antibody staining against desmin in an *A. baerii*. Images show the dorsal branchial region from lateral view. (a``) stage 40. (b``) stage 41. Scale bar is 200 μm.

**Stage 41**(Figure 6b-b``): All *levatores externii* and the *cucullaris* have formed muscle fibers exhibiting intense desmin and 12/101 antibody signals. The *levatores externii* 1 to 4 are all attached to their corresponding epibranchial. The *levator externus* 5 runs ventrad, curving around the epi- and ceratobranchial 4 and tapers into the fibers of the common anlage of the *constrictor pharyngeus* and *transversus ventralis* 5. A few isolated muscle fibers are detectable in the dorsal area of the anlage of the *constrictor pharyngeus*. Ventrally, the *transverus ventralis* 3 has formed muscle fibers exhibiting intense desmin and 12/101 antibody signals. The anlage of the *transversus ventralis* 4 is recognizable as a compact mesenchymal cell cluster ventral to ceratobranchial 4. A muscle anlage, interpreted as the common anlage of the *transversus ventralis* 5 and *coracobranchialis*, emanates from the anlage of the *circumpharyngeus* extending posteroventrad.

**Stage 42**(Figure 7a, b and b`): The *cucullaris* extends more posteroventrad but is still not attached to the supracleithrum. The *transverus ventralis* 4 exhibits intense desmin and 12/101 antibody signals. No desmin and 12/101 antibody signals are detactable in the common anlage of the *circumpharyngeus, transversus ventralis* 5 and *coracobranchialis*.

**Figure 7.**
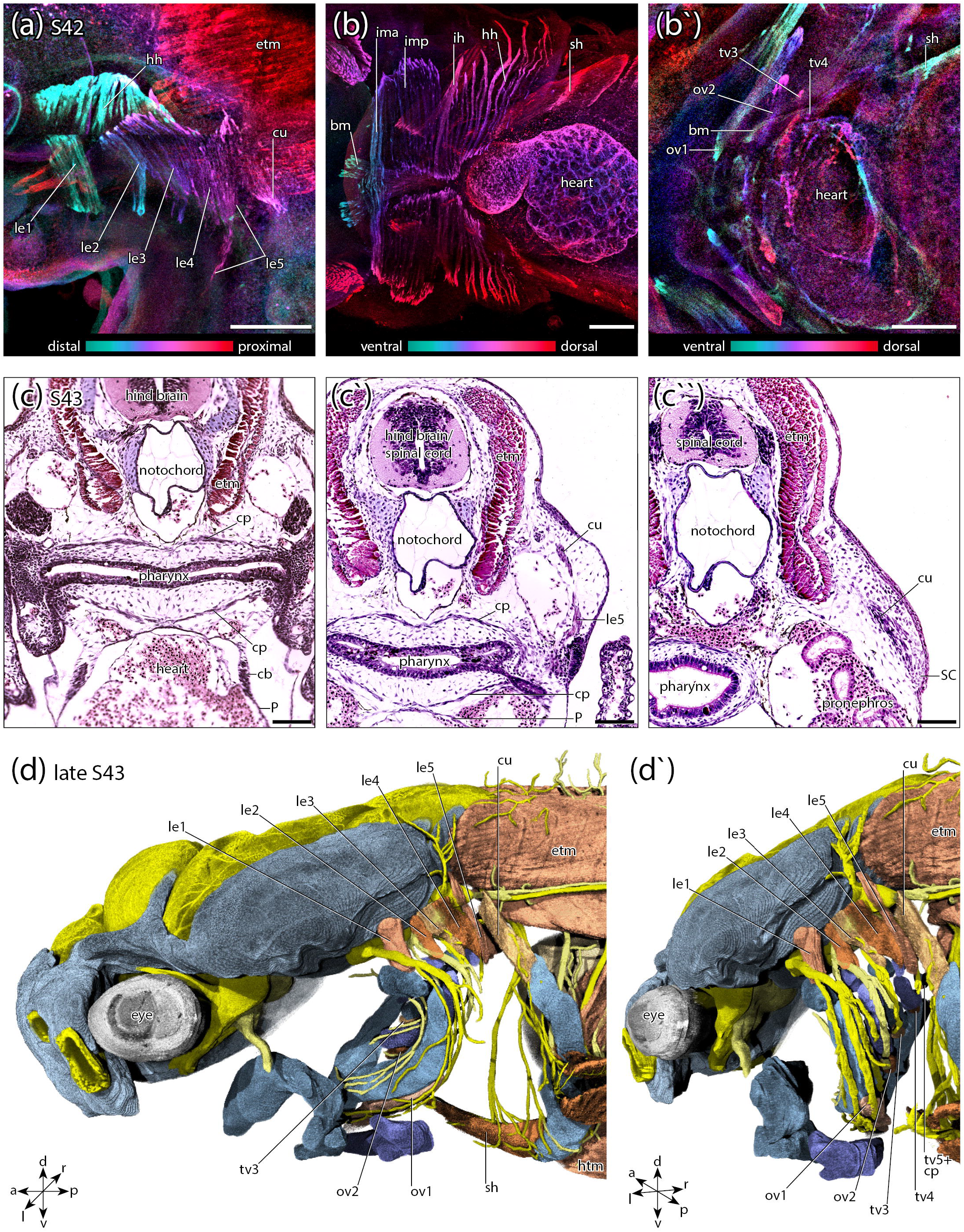
(a to b`) maximum intensity projection of an antibody staining against desmin of an *A. baerii* at stage 42. Colors are depth-coded indicated by the color scale. (a) dorsal branchial region from lateral view. (b) branchial region from ventral view. (b`) branchial region from ventral view. superficial muscles have been removed digitally to show the deeper musculature. (c to c``) histological cross sections through an *A. baerii* at stage 43. (d and d`) 3D reconstruction of the anatomy of the branchial and hypobranchial musculature in *A. baerii* at stage 43. Reconstructions are based on digital image stacks of whole-mount antibody staining against desmin acquired with a laser-scanning microscope. (d) lateral view. (d`) posterolateral view. Scale bar is 200 μm.

**Stage 43**(Figure 7c-d`): The *cucullaris* inserts on the anterior surface of the supracleithrum. The *transversus ventralis* 5 and *circumpharyngeus* are well developed, exhibiting muscle fibers in histological sections. The *coracobranchialis* is more distinct from the *transversus ventralis* 5 now but still not separated from it. It has formed muscle fibers, extends further posteroventrad along the pericard but still not inserts on the anterodorsal edge of the coracoid.

## 4 Discussion

### 4.1 Interspecific variation and the use of model species in evolutionary morphology

Many scenarios on phenotypic evolution of higher-ranked taxa are based on data from only a few model species (Adachi, Pascual-Anaya, Hirai, Higuchi, & Kuratani, 2018; Edgeworth, 1935; Heude et al., 2018; Konstantinidis et al., 2015; Kusakabe et al., 2020; Naumann et al., 2017a; Sefton, Bhullar, Mohaddes, & Hanken, 2016; Stundl et al., 2019; Stundl et al., 2020; Ziermann et al., 2018). This approach, considering limited resources and impeded access to specimens, is regarded suitable if the model species represents a good “cross-section” of morphological and developmental variation of the considered taxon. Nevertheless, variation may occur in other members of the respective taxon, and therefore the information gathered on one species can be misleading, when drawing evolutionary scenarios. One such example is the enigmatic *cucullaris* muscle, which is a complicated case due to the filigree nature of this muscle in many early-branching ray-finned fishes. *Polypterus bichir*, in which a *cucullaris* is reported as absent (Greenwood & Lauder, 1981), would not be an appropriate model to represent the Polypteriformes in questions regarding the reconstruction of the evolution of the vertebrate *cucullaris*, as in *P. ornatipinnes* and *P. senegalus* a *cucullaris* is present (Greenwood & Lauder, 1981; Noda et al., 2017).

The craniofacial skeleton of sturgeons represents a known case of interspecific as well as intraspecific variation (Hilton & Bemis, 1999; Hilton, Dillman, Paraschiv, & Suciu, 2020; Tsessarsky, 2021; Vetter, 1877; Warth et al., 2017). In the present study on the development and morphology of the branchial musculature of sturgeons, we found variation in in comparison to the literature in three distinct cases:

1. *attractores arcuum branchialium* - In *A. baerii*, the *attractores arcuum branchialium* appears to be absent until in specimens smaller than 19.5 cm total length. This is different to larval and juvenile *A. sturio* where the *attractores arcuum branchialium* 1 to 3 (also *adductores arcuum branchialium* see Winterbottom, 1973) are present already at small sizes and to *A. ruthenus* and *A. fulvescens* in which only the *attractor arcuus branchialium* 4 is reported present already at a juvenile stage (Edgeworth, 1929, page 63).
2. *obliquus ventralis* - Edgeworth (1911, page 236) describes a condition in *A. sturio* where the *obliquus ventralis* 2 and 3 insert on the corresponding hypohyals while the *obliquus ventralis* 4 and 5 insert on a median raphe (forming *transversus ventralis* 4 and 5). In a later study (1929, page 73) Edgeworth states, that the *obliquus ventralis* 3 insert on a median raphe (forming *transversus ventralis* 3). Edgeworth is not always precise to which species he refers in his observations (writing mostly “*Acipenser*”). In his study from 1929, Edgeworth reinvestigated *A. sturio* specimens from 1911 (stated in Edgeworth, 1929, page 39). However, he mentioned in the acknowledgements that he received some additional embryos of *A. ruthenus* from Schmalhausen (1929, page 81). Therefore, it is unclear whether he merged his observations made in *A. sturio* with observation made in *A. ruthenus*. This impedes exact inference on interspecific variation of craniofacial muscle development in sturgeons from his works. Our observation on *A. baerii* is consistent with the condition described presumably for *A. sturio* in Edgeworths study from 1929. The ventral muscle of the third branchial arch inserts at a median raphe with its antimere, forming the *transversus ventralis* 3.
3. *Levatores externi, cucullaris and coraco-branchialis* - Regarding the dorsal branchial muscles, different descriptions of the number of *levatores externi* (also *levatores arcuum branchialium*, see Winterbottom, 1972) and the origin and presence of a *cucullaris* (also *protractor pectoralis* and *trapezius*, see Winterbottom, 1972) are available. Four “*levatores arcuum branchialium*” are reported for *A. sturio* by Edgeworth (1911; pages 236-237, Figure 22) with a fifth muscle passing from epibranchial 4 to ceratobranchial 5. Edgeworth mentioned that this muscle might be the *levator arcuum branchialis* 5 described by Vetter (1877) in the same species. Vetter reported five *levatores arcuum branchialium* in *A. sturio*, with the fifth levator inserting on “the posterior-most branchial arch element” (i.e., ceratobranchial 5; Vetter, 1877, page 477). In his 1935 work (page 131), Edgeworth mentioned the presence of five *levatores arcuum branchialium* in “*Acipenser”*, maybe adopting Vetter’s interpretation. However, in his drawing (1935, page 354, Figure 224) he only labeled 3 levator muscles as “Levs. arc. br. II-IV” and located posterior to those, a “cucullaris”. This is consistent with his study from 1911 where he describes the *cucullaris* growing back from the *levator arcuum branchialium* IV (1911, page 236) and hypothesized an origin from the fourth branchial arch muscle plate arch (1911, page 257). In 1935, Edgeworth reported that a *cucullaris* is absent in *A. sturio* but present in *A. ruthenus*, consisting of two portions, “one passing from the cranium to the scapula and cleithrum and the other from the suprascapula to the scapula.” (Edgeworth, 1935, page 142). A *coracobranchialis*, derived from the fourth branchial arch, is absent in *A. sturio* according to Vetter (1874) but has been observed by Edgeworth (1911) in the same species. In *A. ruthenus* it is innervated by a vagal ramus of the fourth branchial arch (Edgeworth, 1929). The *coracobranchialis* of *A. baerii* however develops in contact with the *transversus ventralis* 5 and *constrictor pharyngeus* instead of the *transversus ventralis* 4 implying its origin from the fifth branchial arch in this species. We hypothesize, that many of the different descriptions of the presence/absence of a *cucullaris* in different sturgeon species might be due to the filigree nature of this muscle in many early-branching ray-finned fishes. Whole-mount antibody staining against different muscle proteins has proven detecting also very thin filigreed muscles (Naumann & Olsson, 2018). This technique has been used to clearly identify a *cucullaris* in some early-branching actinopterygian taxa where an absence of this muscle was reported previously (*Lepisosteus osseus*: Naumann et al., 2017 vs. Edgeworth 1935; *Polypterus senegalus*: Noda et al., 2017 vs. Greenwood and Lauder, 1981).

The discussed differences of the development and morphology of the branchial musculature of different *Acipenser* species indicate a possible high degree of interspecific variation in sturgeons. However, it is not clear to what degree observational bias contributes to the described differences in craniofacial muscular morphology. A large-scale comparative study of craniofacial muscle development of several different *Acipenser* species is needed to gain data on the intra- and interspecific variation and clarify this question.

### 4.2 Functional morphology

One of the many biological roles (Bock & von Wahlert, 1965) of the craniofacial system is food uptake. In some non-teleostean actinopterygians (Polypteriformes, Lepisosteiformes), mouth opening is facilitated by two musculoskeletal couplings. The epaxial-neurocranial coupling dosally elevating the head and the coupling involving the hypaxial musculature, the cleithrum and the hyoid- and hypobranchial apparatus ventrally, causing mandibular depression (Liem, Bemis, Walker, & Grande, 2000). This mechanism is also found in early-branching lobe-finned taxa (Dipnoi, Actinistia) and is therefore considered ancestral for osteognathostomes in general and also actinopterygians (Lauder, 1982).

Sturgeons and some chondrichthyans exhibit a distinctive mode of jar suspension in which the palatoquadrate is not fused to the neurocranium but, together with the lower jaw, suspended via a mobile hyoid arch (Bemis et al., 1997; Gegenbaur, 1898). This hyostylic condition evolved independent within the two lineages and kinematic studies have been carried out in some chondrichthyan species (Ferry-Graham, 1997; Frazzetta & Prange, 1987; C Wilga & Motta, 1998) and *Acipenser medirostris* (Carroll & Wainwright, 2003) to resolve the functional morphology of this anatomical condition. It has been hypothesized that hyoid (= anterior and posterior ceratohyals and hypohyals) retraction is mediated by contraction of the *sternohyoideus* in *Scaphirhynchus albus* and *A. medirostris* (Carroll & Wainwright, 2003). In the study of *S. albus* by Carroll and Wainwright (2003), only the “primary” insertion site of the *sternohyoideus* on the hypohyal is mentioned. Our investigation of the branchial and hypobranchial musculature of *A. baerii* revealed three insertion sites of the *sternohyoideus* (hypohyals, 1. and 2. hypobranchials) and a switch of the origin of the *obliquus ventralis* 1 from hypobranchial 1 to the hypohyal and the anterior ceratohyal. This is consistent with reports for *A. sturio* and possibly *A. ruthenus* (Edgeworth, 1911, 1929, 1935). This musculoskeletal configuration suggests a more complex feeding mechanism, involving branchial basket retraction via the strong *obliquus ventralis* 1 and *sternohyoideus* parallel to lower jaw depression and partial protrusion. Kinematic studies involving marker based x-ray motion analysis (Camp & Brainerd, 2015; Schwarz, Gorb, Kovalev, Konow, & Heiss, 2020) would be helpful to reevaluate sturgeon feeding kinematics and unravel the potential role of the *obliquus ventralis* 1 in jaw opening.

### 4.2 The role of heterochrony in sturgeon craniofacial muscle development

In the present study, we show that *A. baerii* exhibits a distinctive development and morphology of some craniofacial muscles. The anteriormost branchial muscles, the *levator externus* 1 and *obliquus ventralis* 1, develop early (starting during late stage 36) compared to the more posterior branchial muscles starting at stage 39. Additonally, *A. baerii* and other sturgeons exhibit an unusual morphology of the *sternohyoideus*, inserting not only on the hypohyal but also on hypobranchial 1 and 2, and the *obliquus ventralis* 1, originating from the skeleton of the hyoid arch (the hypohyal and anterior ceratohyal) instead of the skeleton of the first branchial arch. In contrast, the late developmental timing of the posterior-most branchial muscles (e.g. the *cucullaris* and the *coracobranchialis*) differentiating at the head-trunk interface is similar to some other osteognathostomes (Diogo et al., 2008; Edgeworth, 1935; Ericsson, Knight, & Johanson, 2013; Naumann & Olsson, 2018; Naumann et al., 2017a; Noda et al., 2017; Sefton et al., 2016; Theis, 2010; Ziermann et al., 2018; Ziermann & Diogo, 2013). The cellular mechanism underlying the peculiar morphology (hyoid association of the *obliquus ventralis* 1 and branchial association of the *sternohyoideus*) and the heterochronic shift of some branchial muscles (precocious or accelerated development of the *levator externus* 1 and *obliquus ventralis* 1) of *A. baerii* is still unclear. An attempt to explain this condition is based on the migratory pattern of the cranial neural crest cells observed in *A. ruthenus* (Stundl et al., 2020). In this species, the development of the otic capsule is delayed resulting in an undivided hyobranchial neural crest sheet emigrating from the neural tube. Later, the developing otic capsule and second pharyngeal pouch result in a postponed separation of the hyoid and anterior branchial neural crest streams (Stundl et al., 2020). This might result in an early interaction of hyoid/branchial neural crest cells with mesodermal muscle precursors of the first branchial arch accelerating their differentiation. Furthermore, this incomplete division of the hyoid and branchial neural crest might result in the unusual association of the *sternohyoideus* with branchial and the *obliquus ventralis* 1 with hyoid skeletal elements. While this hypothesis is rather speculative it can be supported with data on the derived developmental pattern of endo- and mesodermal structures similarly correlating with a heterochronic shift of the migration of the associated neural crest cells that has been reported for *Polypterus senegalus* and *Lepisosteus osseus* (Stundl et al., 2019; Stundl et al., 2020).

### 4.4 Propagation-based phase contrast tomography

A variety of techniques is available for studies of developmental morphology and with technological advances, new options arise. In here we combined classical histology, whole-mount antibody staining and propagation-based phase contrast tomography. This technique allows producing high-resolution digital image stacks from iodine-stained samples within several minutes and the combination with robotics therefore enables high-throughput workflows. The resolution is sufficient to clearly identify a variety of anatomical structures (Figure 7a to d) and subsequently prepare high quality 3D reconstructions of early embryos (Figure 7e to f```). Especially for early- to mid-embryonic stages, this technique was favourable and yielded additional insights not evident from the fluorescent antibody stainings investigated by laser scanning microscopy. Yet the combination with fluorescent antibody stainings as well as histology allowed detailed comparison and confirmation of tissue identity where selective stainings revealed additional information. On the other hand, especially early embryonic stages are difficult to section and histological slides therefore sometimes suffer from deformation and breakage. Hence 3D reconstructions from histological slides are not always possible and at least very time consuming, making propagation-based phase contrast tomography a valuable technique to study and display whole-mount embryos.

## Acknowledgements

We would like to thank Katja Felbel for help with histological sections. Peter Groß and his team at the Fischzucht Rhönforelle kindly supplied us with fertilized eggs of *A. baerii*. The monoclonal antibodies obtained from the Developmental Studies Hybridoma Bank were developed under the auspices of the NICHD and maintained by The University of Iowa, Department of Biological Sciences, Iowa City, IA 52242, USA. PW received a scholarship from the Landesgraduiertenförderung Thüringen. We acknowledge DESY (Hamburg, Germany), a member of the Helmholtz Association HGF, for the provision of experimental facilities. Parts of this research were carried out at PETRA III during beam times related to the proposals I-20140659 to JUH and I-20160119 to BN. This research was supported in part through the Maxwell computational resources operated at Deutsches Elektronen-Synchrotron (DESY), Hamburg, Germany.

## Abbreviations

a: anterior
aCH: anterior ceratohyale
am: adductor mandibulae
BC: basibranchial copula
BH: basihyale
bm: branchiomandibularis
cb: coracobranchialis
CB1-5: ceratobranchial 1 to 5
CH: ceratohyale
CO: coracoid
cp: constrictor pharyngeus
cu: cucullaris
d: dorsal
e: eye
EB1-4: epibranchial 1 to 4
etm: epaxonic trunk musculature
fm: pectoral fin musculature
gPLLN: ganglion of the posterior lateral line nerve
gIX: ganglion of the glossopharyngeal nerve
gX_b2_: anterior vagal ganglion
gX_b3-b5_: posterior vagal ganglion
HAA: hyoid arch artery
HB1-3: hypobranchial 1 to 3
hh: hyohyoideus
HH: hypohyale
HM: hyomandibula
htm: hypaxonic trunk musculature
ih: interhyoideus
IH: interhyale
im: intermandibularis
IX_le1_: branch of the glossopharyngeal nerve innervating the levator externus 1
l: left
MC: Meckel’s cartilage
NA: nasal
le1-5: levator externus 1 to 5
NC: notochord
OC: occipital
OT: otic capsule
ov1-2: obliquus ventralis 1 to 2
p: posterior
P: pericard
PP: parachordal plate
pCH: posterior ceratohyale
ph: protractor hyoideus
PH: pharynx
pNW: posterior nasal wall
PQ: palatoquadrate
r: right
rh: retractor hyoideus
S2_sh_: branches of spinal nerve 2 that innervate the sternohyoideus in addition to the hypoglossal nerve
SC: scapula
sh: sternohyoideus
tcu: tendon of the cucullaris muscle
tv3-5: tranversus ventralis 3 to 5
v: ventral
X_cu_: branch of the vagal nerve innervating the cucullaris
X_le1-le4_: branch of the vagal nerve innervating the corresponding levator externus; levator externus 5 is innervated by a nerve originating from X_le4_
X_ov1-ov2_: branch of the vagal nerve innervating the corresponding obliquus ventralis
X_tv3-4_: branch of the vagal nerve innervating the corresponding transversus ventralis
X_tv5+cp_: branch of the vagal nerve innervating the not completely separated transversus ventralis 5 and constrictor pharyngeus
XII_bm_: branch of the hypobranchial nerve innervating the branchiomandibularis
XII_sh_: branch of the hypobranchial nerve innervating the sternohyoideus

**Supplementary Material 1**. Complete list of specimens. AB, whole-mount antibody staining; IACT, iodine-enhanced absorption contrast tomography; PPCT, propagation-based phase contrast tomography; Re-Tri, Resorcin-Trichrome staining (histology).

**Supplementary Material 2**. 3D reconstruction of craniofacial skeletal elements. Cartilaginous elements are colored in different shades of blue. The notochord is colored in ivory. Naming of skeletal elements is according to Hilton, Grande and Bemis, 2011. (a) head and shoulder girdle from lateral view. (b to d) isolated branchial basket. (b) ventral view. (c) lateral view. (d) dorsal view.

