## Supplementary figures and images for "Development of the branchial musculature of the Siberian sturgeon (*Acipenser baerii*) reveals a heterochronic shift during the evolution of acipenseriform cranial muscles"

### Supplementary Material 1

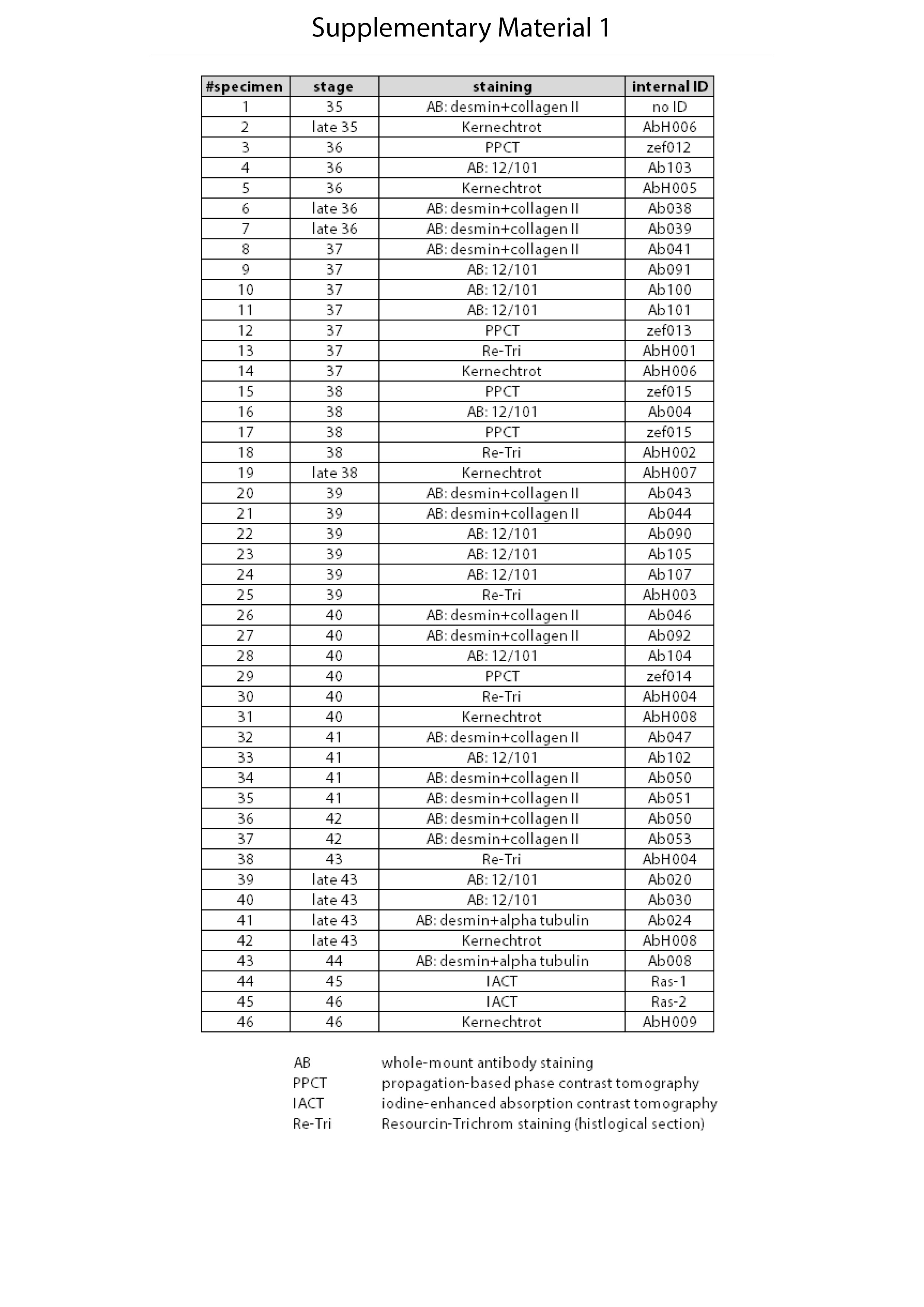

### Supplementary Material 2

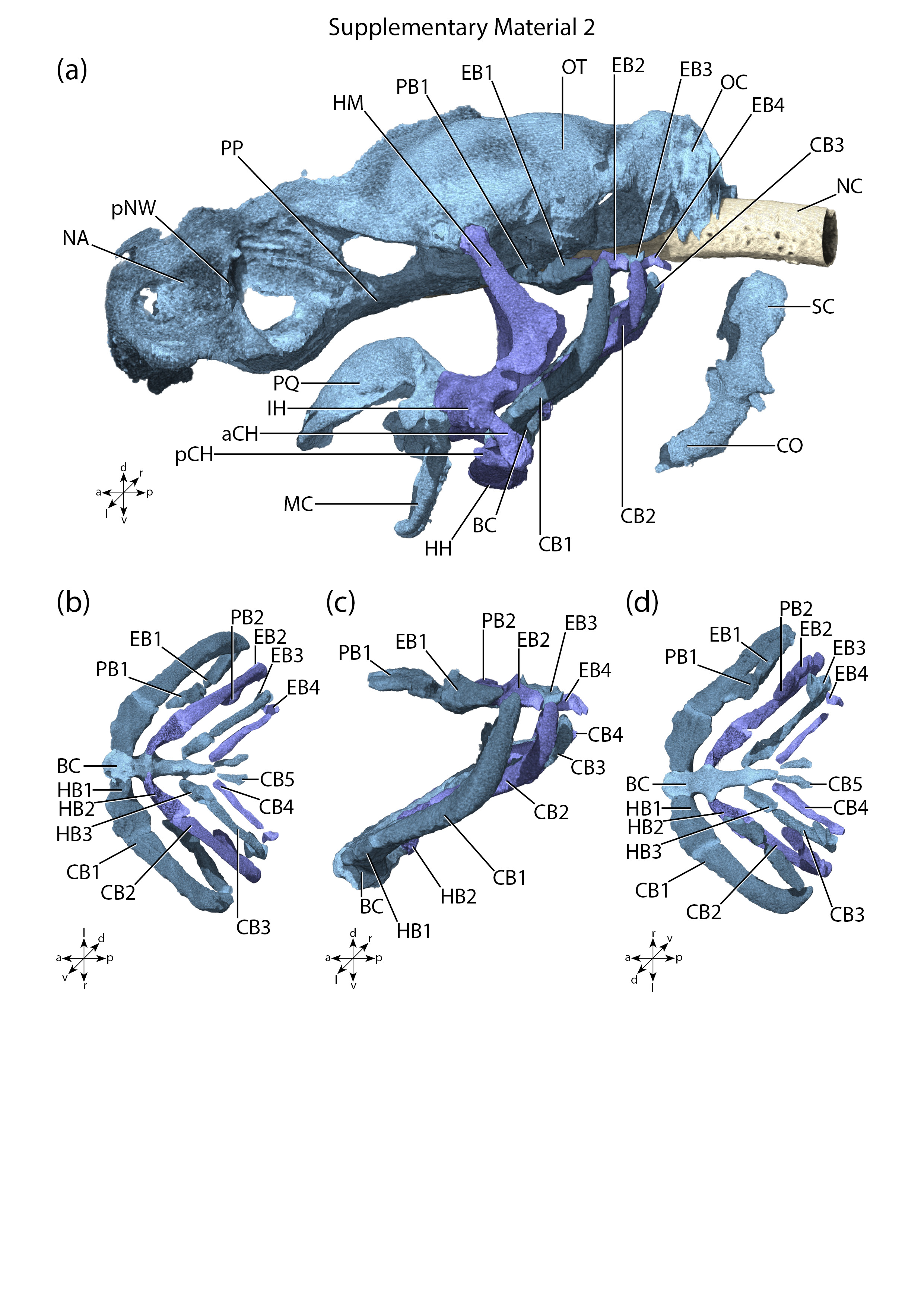
